# Neurotrophin-3 produced by motor neurons non-cell autonomously regulates the development of pre-motor interneurons in the developing spinal cord

**DOI:** 10.1101/2025.07.27.665825

**Authors:** Andrea Angla-Navarro, Ana Dominguez Bajo, Mathilde Toch, Cédric Francius, Maria Hidalgo-Figueroa, Jingwen Zhang, Mélina Finet, Olivier Schakman, Manon Martin, Xiuqian Mu, René Rezsohazy, Françoise Gofflot, Frédéric Clotman

## Abstract

The development of multicellular organisms requires proper interplays between cell-autonomous genetic programs controlled by combinations of transcription factors that regulate the differentiation of distinct cell populations and non-cell autonomous processes that coordinate the proliferation, the fate, the survival, the respective location, and the proper interactions of these populations. During the development of the nervous system, non-cell autonomous mechanisms determine neuronal fate, survival, distribution, axon guidance, and connectivity. Although similar processes are suggested to be at work in the formation of spinal motor circuits, the molecular mechanisms involved remain mostly elusive. Here, we provide evidence that the Onecut transcription factors regulate a non-cell autonomous mechanism that modulate pre-motor interneuron development. We show that conditional inactivation of the Onecut factors in spinal motor neurons affects the differentiation and the positioning of pre-motor interneuron populations. We identify that Neurotrophin-3 produced by motor neurons under the control of the Onecut factors non-cell autonomously regulate the production and the distribution of pre-motor interneuron populations. Thus, we elucidated one of the non-cell autonomous mechanisms that coordinate the formation of the spinal motor circuits.

## INTRODUCTION

The development of multicellular organisms requires the coordinated production and positioning of a constellation of specialized cells exerting distinct but complementary functions. Specialization of stem or progenitor cells into different cell types depends on the activation of distinct cell-autonomous genetic programs that are controlled by time- and space-regulated combinations of transcription factors (Allan and Thor, 2015; Sagner, 2024). However, this intrinsic regulation additionally necessitates extrinsic mechanisms that coordinate the proliferation, the fate, the survival, the respective location, and the proper interactions of these different cell populations (Czesnick and Lenhard, 2015; Surya and Sarinay-Cenik, 2022; Williamson, 2023). These non-cell autonomous processes, which rely on signals coming from the extracellular environment including secreted molecules, cell-cell interaction through membrane receptors, changes in electrical activity or mechanical forces (see for example (Choi et al., 2020; Guemez-Gamboa et al., 2014; Lim et al., 2018; Pillai and Franze, 2024; Welniarz et al., 2017)), have been less extensively studied. Nevertheless, the emergent picture reveals a dynamic landscape where signaling pathways, diffusible factors, cell activity and mechanical forces traverse cell membranes, enabling cells to communicate, cooperate and adapt, to generate functional multicellular tissues or organs.

During nervous system development, non-cell autonomous mechanisms determine neuronal fate, survival, distribution, axon guidance, and connectivity. For example, Cajal-Retzius (CR) cells, a transient class of neurons that populate the mantle zone of the mammalian cerebral cortex very early during brain development, regulate the respective size of different cortical areas (Barber et al., 2015), critically control the laminar organization of the neocortex through the secretion of Reelin (reviewed in (Jossin, 2020)) and of Nectin1 (Gil-Sanz et al., 2013), and their activity alters survival and connectivity of pyramidal neurons (Riva et al., 2019). Inversely, CR cell migration is controlled by an extracellular contingent of the homeoprotein Pax6 secreted by cortical progenitors (Kaddour et al., 2020). The proliferation, survival and radial growth of cortical progenitors and neurons are regulated by neurons that originate outside of the neocortex (Causeret et al., 2011; Teissier et al., 2010). Reciprocally, the identity, connectivity and survival of cortical interneurons (INs), which are born in subcortical ganglionic eminences and colonize the neocortex during development, is determined by the type of cortical projection neurons they approach (Lim et al., 2018; Wester et al., 2019), and their number and organization in the marginal zone are controlled by non-muscle myosin II heavy chain-mediated regulation of radial glial cell complexity and endfoot position (D’Arcy et al., 2023). Among cortical neurons, the transcription factor Foxp2 produced in layers 5 and 6 of the cerebral cortex regulate in a non-cell autonomous manner gene expression in projection neurons and the number of cortical INs (Co et al., 2020). The distribution of cortical INs is also modulated by extrinsic signaling. The proto-cadherin PCDH19 non-cell autonomously affects cell migration in medial ganglionic eminence explants (Pancho et al., 2022), and netrin signaling determines the migration of cortical INs (Yamagishi et al., 2020). In addition, axonal growth and neurite formation are modulated by extrinsic processes. Cunningham and colleagues (Cunningham et al., 2022) demonstrated that the complete genetic removal of the c-Jun N-terminal kinase gene, which is expressed in striatal, guidepost and corridor cells, causes the inability of thalamocortical axons to cross the diencephalon-telencephalon boundary, causing axonal misrouting to other cerebral areas. The netrin receptor DCC controls midline crossing of the cortico-spinal tract in a non-cell autonomous manner in mice, but not in humans (Welniarz et al., 2017), whereas lumican, an extracellular proteoglycan secreted by corticospinal lateral neuronal populations, extrinsically suppresses cervical collateralization by multiple corticospinal medial subpopulations (Itoh et al., 2023). Inversely, the ascending axonal growth of rapidly adapting mechanoreceptors to the brainstem is modulated by roof plate radial glial-like cells (Kridsada et al., 2018). Neuronal activity also triggers non-cell autonomous processes. Activity-dependent BDNF release regulates the Glutamate-GABA neurotransmitter switching in a non-cell autonomous manner in Xenopus (Guemez-Gamboa et al., 2014), and the identity and position of dorsal root ganglia neurons in mice (Wright and Ribera, 2010). In zebrafish, voltage-gated Na+ channels expressed in specific motor neuron (MN) subtypes regulate axonal projections of other MNs that do not produce them (Pineda et al., 2006), and in the mouse GluR1-dependant activity regulates MN neurite formation and connectivity in a cell autonomous and a non-cell autonomous manner (Zhang et al., 2008). Finally, terminal maturation of neural circuit is coordinated by extrinsic cues. For example, critical periods of heightened IN plasticity are defined by multiple non-cell autonomous signals, such as brain-derived neurotrophic factor (BDNF), the synapse-regulating SPARCL1 glycoprotein, the homeoprotein OTX2 or extracellular matrix components, produced by cortical or extra-cortical sources (Bernard et al., 2016; Gibel-Russo et al., 2022; Vincent et al., 2021).

Furthermore, different pieces of published data suggest that similar coordination mechanisms are also at work in the developing spinal cord. Body movements are regulated by neural circuits located in ventral regions of the spinal cord and constituted of MNs, which directly innervate skeletal muscles, and of multiple populations of pre-motor INs, the integrated actions of which control the activity of the MNs (Cote et al., 2018). Some aspects of spinal motor circuit development are reported to rely on non-cell autonomous mechanisms. Differentiating MNs modulate the entry of MN progenitors into differentiation through a regulation of the Notch pathway involving the six-transmembrane protein glycerophosphodiester phosphodiesterase 2 (GDE2) and disinhibition of the ADAM protease by release of the RECK activator from the MN membrane (Park et al., 2013; Sabharwal et al., 2011). During V2 IN differentiation, the transcription factor GATA2, which is present in progenitors and in differentiating V2 cells, regulate in a non-cell autonomous manner the proliferation of neural progenitors through an unknown mechanism (El Wakil et al., 2006). In zebrafish, early-born primary MN determine the GABAergic phenotype of Kolmer–Agduhr INs (Seredick et al., 2014). Constitutive inactivation of *Islet-1* (*Isl1*), a transcription factor present in MNs and in dorsal dI3 INs, alters the development of V1 INs (Pfaff et al., 1996). Similarly, the transcriptional regulator Bhlhb5 influences the generation of INs surrounding its distribution domain (Skaggs et al., 2011), and inactivation of WT1 from dI6 INs changes the composition in V0 and in V2a INs (Schnerwitzki et al., 2018). Furthermore, the survival of spinal IN populations is controlled by extrinsic cues, as their apoptosis is differentially regulated by protocadherin-gamma adhesion molecules (Prasad et al., 2008). Taken together, these observations demonstrate that, in the nervous system, non-cell autonomous mechanisms regulate neuronal fate, survival, distribution, and connectivity. However, how much similar processes contribute to coordinate the development of neuronal populations forming the spinal motor circuits remains unknown.

Onecut (OC) transcription factors, including Onecut-1 (OC-1), OC-2 and OC-3, are homeodomain-containing transcriptional regulators that control multiple aspects of motor circuit development, including MN or IN differentiation or survival (Francius and Clotman, 2010, 2014; Roy et al., 2012; Stam et al., 2012; Toch and Clotman, 2019; Toch et al., 2020), IN distribution (Harris et al., 2019; Kabayiza et al., 2017), and formation of the neuromuscular junctions (Audouard et al., 2012). Here, we provide evidence that the OC factors regulate in spinal MN a non-cell autonomous mechanism that modulate pre-motor IN development. We identify candidate factors potentially involved in this process, and we demonstrate that Neurotrophin-3 (Ntf3) produced by MN contributes to the regulation of IN differentiation and distribution, and thereby coordinate proper formation of spinal motor circuits.

## RESULTS

### OC factors regulate in spinal MNs a non-cell autonomous mechanism that modulates pre-motor IN development

The OC transcription factors are produced in multiple spinal neuronal populations including MNs (Francius and Clotman, 2010; Francius et al., 2013; Roy et al., 2012) and ventral (Francius et al., 2013; Harris et al., 2019; Stam et al., 2012; Toch et al., 2020) or dorsal INs (Kabayiza et al., 2017). However, OC are not present in all the cells of each population but are produced in specific subsets (Francius et al., 2013; Harris et al., 2019; Stam et al., 2012). In V1 INs, OC are detected in Renshaw cells (RC) but in none of the other V1 components (Francius et al., 2013; Stam et al., 2012). In the course of our studies investigating the function of the OC proteins during spinal cord development in constitutive *Oc1/Oc2* double-mutant embryos (Harris et al., 2019; Stam et al., 2012; Toch et al., 2020), which additionally lack OC-3 in the spinal cord (Kabayiza et al., 2017; Roy et al., 2012), we noticed alterations in the differentiation of non-RC V1 INs (Figure 1A-C). In particular, a subset of Foxp2+ V1 INs containing the MafA transcriptional regulator increased in cell number and expanded in an abnormal ventro-medial location as compared to control embryos at embryonic day (e)11.5 (Figure 1A, C). Similarly, V1 INs abnormally producing Prdm8 (Francius et al., 2013) and Otp were observed in the same ventro-medial position at e12.5 (Figure B-C). These observations suggested that OC factors may control in the developing spinal cord a non-cell autonomous mechanism regulating the differentiation of V1 INs.

**Figure 1.**
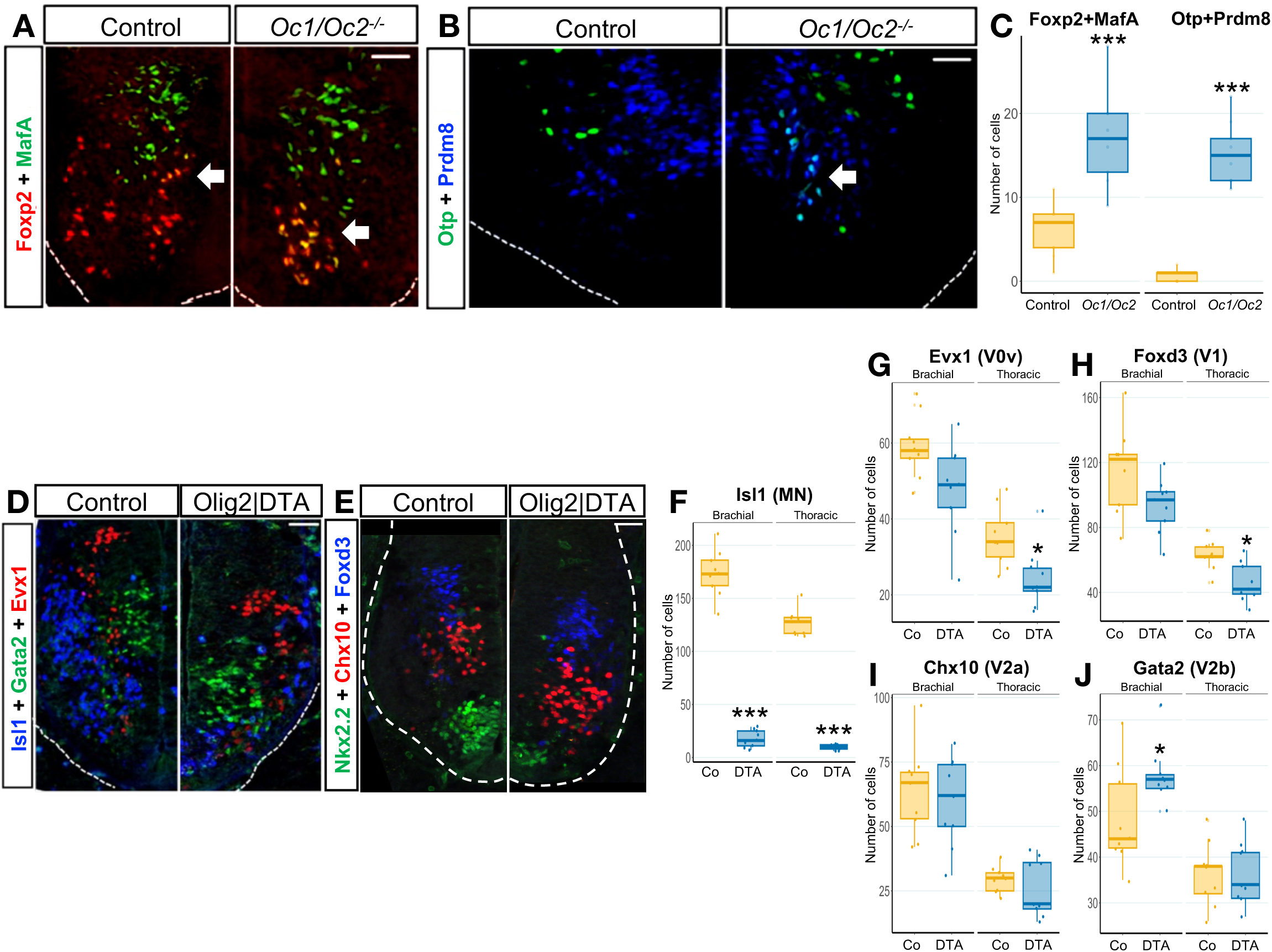
Spinal interneuron development is altered in Onecut constitutive mutants and in embryos depleted in spinal motor neurons. (A-C) In constitutive *Oc1/Oc2* double-mutant embryos, which additionally lack OC-3 in the spinal cord, a subset of Foxp2+ V1 INs containing the MafA transcriptional regulator (yellow) increased in cell number and expanded in an abnormal ventro-medial location as compared to control embryos at embryonic day (e)11.5 (arrows; control left hemicord, mutant right hemicord; thoracic levels). Similarly, V1 INs abnormally producing Prdm8 and Otp (cyan) were observed in the same ventro-medial position at e12.5 (arrow). (D-J) After genetic ablation of ∼90% of spinal MNs, the number of V0v INs characterized by the presence of Evx1, and of V1 INs assessed by the production of Foxd3, was significantly reduced at thoracic levels of the spinal cord. In contrast, the number of V2b INs, which contain Gata3, was increased at brachial levels, whereas V2a INs identified by the presence of Chx10 were unchanged. Scale bars=100μm. n=3; *=adjusted p value < 0.05; ***=adjusted p value < 0.001

Given that OC proteins are critical for proper differentiation of the spinal MNs and expression of *Isl1* in these cells (Roy et al., 2012), that IN alterations have been reported in embryos wherein MN development is strongly affected by the absence of Isl1 (Pfaff et al., 1996), and that MNs exert non-cell autonomous regulation of the differentiation of surrounding cells (Park et al., 2013; Sabharwal et al., 2011), we hypothesized that OC factors may control this non-cell autonomous mechanism by acting in spinal MNs. To assess the possible contribution of MNs to this mechanism, we first investigated the development of pre-motor IN populations after genetic ablation of spinal MNs. Embryos wherein diphteria toxin subunit A (DTA) is produced specifically in MN progenitors upon *Olig2-Cre* activation (Dessaud et al., 2007) showed a >90% reduction in the number of differentiating MN at e11.5 (Figure 1D,F), without alteration of the pMN progenitor domain (Supplemental Figure S1). In these embryos, the number of V2b INs labelled for Gata2 was increased at brachial levels of the spinal cord (Figure 1D-E,J). In contrast, the amount of V0v characterized by the presence of Evx1 and of V1 assessed by the production of Foxd3 (Francius et al., 2013) was significantly reduced at thoracic levels (Figure 1D-E,G-H), while the production of V2a INs was not changed (Figure 1D-E,I). The V3 INs were not investigated, as the *Olig2-Cre* allele also drives recombination, and therefore DTA production, in this population (Chen et al., 2011; Debrulle et al., 2020). Distribution of the pre-motor IN population was not analyzed in detail, as the ablation of MNs resulted in a global ventral displacement of all the IN populations (Figure 1D-E). Nevertheless, these observations suggest that MNs modulate the development of surrounding pre-motor IN populations in the developing spinal cord.

To confirm this interpretation and the possible implication of OC factors in this process, we studied the phenotype of ventral spinal INs after combined conditional inactivation of *Oc1* and *Oc2* in spinal MNs at e12.5 (Figure 2) and at e14.5 (Supplemental Figure S2)(Toch et al., 2020). Absence of OC factors does not alter the number of MNs(Roy et al., 2012). The total number of V1 INs, assessed here by the presence of Foxp2 (Francius et al., 2013), of V2a and of V2b INs was not modified when OC factors were absent from MNs (Figure 2A-F; Supplemental Figure S2A-C). In contrast, the number of V2c INs, which contain Sox1 and are located outside of the ventricular zone in the vicinity of the MNs (Panayi et al., 2010), was drastically reduced in the conditional *Oc* double-mutant embryos (Figure 2G-H; Supplemental Figure S2D), without any increase in spinal cord cell death (Roy et al., 2012). Thus, except for the V2c population, absence of OC factors from the MNs does not alter the global number of ventral INs, consistent with previous observations in the constitutive OC mutants wherein the size of the IN populations was not altered (Harris et al., 2019; Toch et al., 2020).

**Figure 2.**
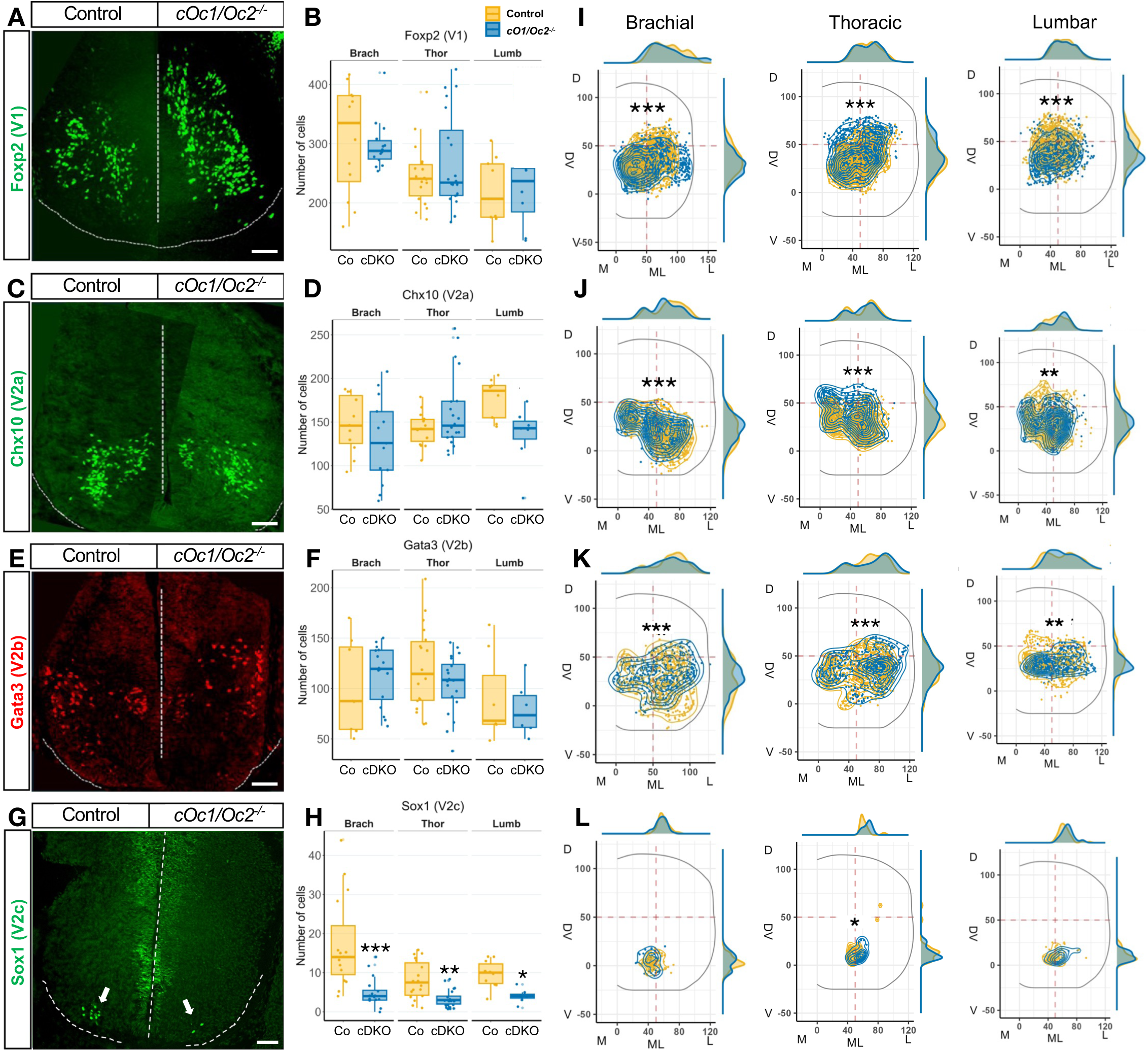
Spinal interneuron number and positioning is altered when Onecut factors are absent from the spinal motor neurons. **(A-L)** In MN conditional *Oc1/Oc2* double-mutant embryos (cDKO) at e12.5, the number of V2c INs identified by the presence of Sox1 outside of the ventricular zone was significantly reduced at all levels of the spinal cord. Furthermore, the positioning of the V1 (Foxp2), V2a (Chx10), V2b (Gata3) and V2c INs on the transversal plane of the spinal cord was significantly altered. Labelings are shown at thoracic levels of the spinal cord. Scale bars=100μm. ML: medio-lateral axis; DV: dorso-ventral axis. n=3; *=adjusted p value < 0.05; **=adjusted p value < 0.01; ***=adjusted p value < 0.001

In contrast, the distribution of ventral IN populations on the transverse plane of the spinal cord was systematically modified, as observed in constitutive OC mutants (Harris et al., 2019; Kabayiza et al., 2017; Toch et al., 2020). In particular, the V1 population was slightly but significantly dorsally mislocated at thoracic levels and ventrally displaced at brachial and lumbar levels in the conditional OC double-mutant embryos at e12.5, as compared to control embryos (Figure 2I). At e14.5, V1 INs settled slightly more dorsal at brachial and lumbar levels while they migrated more ventrally in the control embryos (Supplemental Figure S2E). At e12.5, V2a INs were more dorso-medially located at brachial and thoracic levels but remained more medial at lumbar levels at e12.5 (Figure 2J), whereas at e14.5 they settled more dorsal at brachial and thoracic levels and more ventral at lumbar levels (Supplemental Figure S2F). In contrast, V2b INs were less scattered than in the control embryos, concentrating in a more medial position along the dorso-ventral axis of the spinal cord, and more laterally at thoracic and lumbar levels at e12.5 (Figure 2K) and at e14.5 (Supplemental Figure S2G). Finally, although the population was smaller than in controls, some V2c INs were located more laterally at thoracic levels at e12.5 (Figure 2L) but retained this position while control V2c migrated more laterally at e14.5 (Supplemental Figure S2H). Taken together, these observations indicate that OC factors regulate a non-cell autonomous mechanism in MNs that significantly impact the distribution of pre-motor IN populations in the developing ventral spinal cord.

### OC factors moderate the expression of *Neurotrophin-3* (*Ntf3*) in spinal MNs

To identify genes downstream of OC factors in spinal MNs that could contribute to this non-cell autonomous mechanism, we conducted a bulk RNA-sequencing comparison of the transcriptome of control and of OC-deficient MNs at e10.5 (Figure 3A; GEO repository, accession number: GSE141949)(Toch et al., 2020). This transcriptomic analysis unveiled changes in the expression levels of genes coding for membrane proteins or secreted factors, potentially involved in non-cell autonomous processes (Figure 3B-C; Supplemental Figure S3). Among these, the expression of *Frem1* and *2* (Fras1-related extracellular matrix proteins), *Fat3* (FAT atypical cadherin 3), *Sema3d* encoding a secreted signaling protein of the Semaphorin III family and the neurotrophin receptor *Gfra3* (GDNF family receptor alpha 3) was downregulated in the absence of OC factors. In contrast, expression of *Tac1* (Tachykinin precursor 1) encoding four products of the tachykinin peptide hormone family (substance P, neurokinin A, and neuropeptides K and gamma) that can act as neurotransmitters, *Smim18* (Small integral membrane protein 18), *Cbln2* coding for the secreted glycoprotein Cerebellin-2 and the neurotrophic factor *Ntf3* was increased (Figure 3D and data not shown). Possible impact of MN expression of these genes on the development of pre-motor INs was assessed by overexpression or downregulation after chicken embryonic spinal cord electroporation. However, only *Ntf3* overexpression did mildly alter ventral IN development (Supplemental Figure S4 and data not shown). To assess the expression pattern of *Ntf3* in the developing spinal cord and confirm increased expression in the absence of OC factors, we conducted *in situ* hybridization for *Ntf3* on control or conditional OC double-mutant spinal cord sections at e12.5 (Figure 3D). In control embryos, a weak signal for *Ntf3* was detected in the ventral spinal cord at the location of the motor columns (Figure 3D), consistent with the reported expression of *Ntf3* specific to spinal MNs that decreases between e10.5 and e13.5(Buck et al., 2000; Henderson et al., 1993; Usui et al., 2012). In conditional mutant embryos, the expression level of *Ntf3* was increased and became readily observable but remained restricted to the MNs of the spinal cord (Figure 3D). This indicates that Ntf3 is normally produced by MNs in the developing spinal cord and that OC factors moderate *Ntf3* expression levels in these cells. To determine which cells could respond to Ntf3 secretion by MNs, we performed immunodetection of its cognate receptor TrkC. TrkC was broadly detected in the ventral half of the spinal cord including V1, V2a and V2b interneurons, except in newly-born medial cells (Figure 3E-G), indicating that it is present at the membrane of differentiating ventral INs (Henderson et al., 1993; Usui et al., 2012). Thus, Ntf3 could participate in the non-cell autonomous control of pre-motor IN development by the spinal MNs.

**Figure 3.**
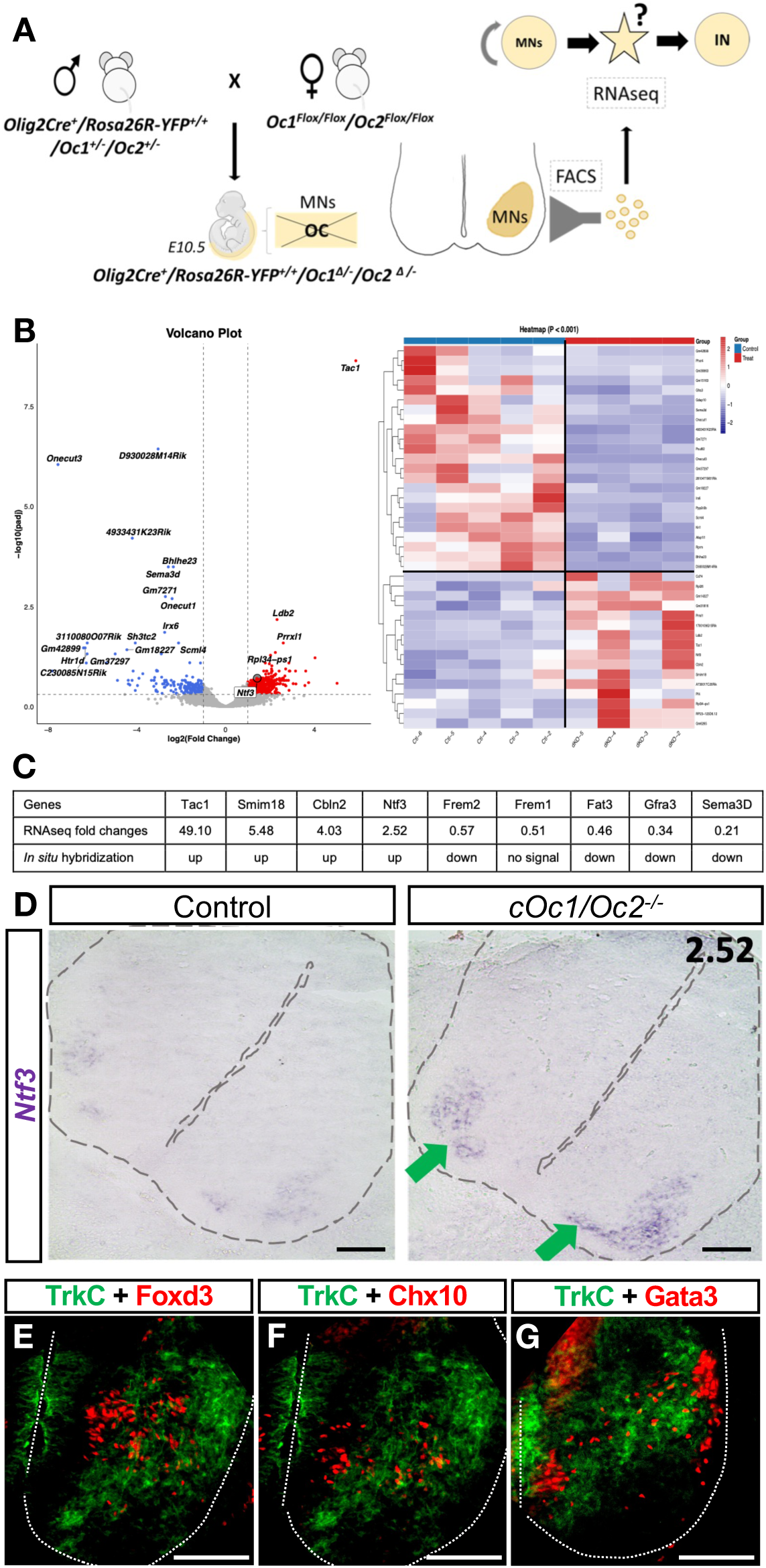
Ntf3 is expressed in spinal motor neurons under the control of the Onecut transcription factors. **(A)** RNA-sequencing comparison of the transcriptome of control and of OC-deficient MNs at e10.5. The spinal cord of control or MN conditional *Oc1/Oc2* double-mutant embryos was isolated, dissociated, YFP-positive MNs were FACS-sorted, RNA was extracted and compared by bulk RNA sequencing. **(B)** Volcano plot or heat-map of the most down- or up-regulated genes in spinal MN deficient in Onecut factors. *Ntf3* is highlighted in the Volcano plot (see GEO repository, accession number GSE141949 for more details). **(C)** Transcriptomic analysis unveiled changes in the expression levels of selected genes encoding membrane proteins or secreted factors. MN-specific expression and changes in expression levels were confirmed by *in situ* hybridization (n=5). **(D)** Expression of the neurotrophic factor *Ntf3* was specific to MNs and increased in the absence of Onecut factors (green arrows). Labelings are shown at thoracic levels of the spinal cord. **(E-G)** TrkC, the cognate receptor of Ntf3, was broadly detected in the ventral half of the spinal cord including V1 (Foxd3, E), V2a (Chx10, F) and V2b (Gata3, G) interneurons, with the exception of newly-born neurons located in medial position. Scale bars=100μm.

### Ntf3 production by MNs is required for proper development of pre-motor INs

To validate this hypothesis, we conditionally inactivated *Ntf3* in spinal MNs and analyzed the production and distribution of pre-motor IN populations. *Ntf3* inactivation did not increase spinal cord cell death (Supplemental Figure S5). In embryos carrying a MN-specific deletion of *Ntf3* at e12.5, the number of V1 and of V2a INs was significantly increased at all levels of the spinal cord (Figure 4A-D). Similarly, a significant increase in V2b was observed at thoracic levels (Figure 4E-F), whereas V2c INs were unchanged (Figure 4G-H). At e14.5, the number of V1 was still increased at all levels (Supplemental Figure S6A), whereas V2a increase was only maintained at lumbar levels (Supplemental Figure S6B), suggesting that programmed cell death of INs that is initiated at this stage (Prasad et al., 2008) may be increased in the absence of Ntf3. Nevertheless, impaired production of Ntf3 by the MNs results in an excess of spinal pre-motor INs in the V1 and V2 populations (Supplemental Figure S6).

**Figure 4.**
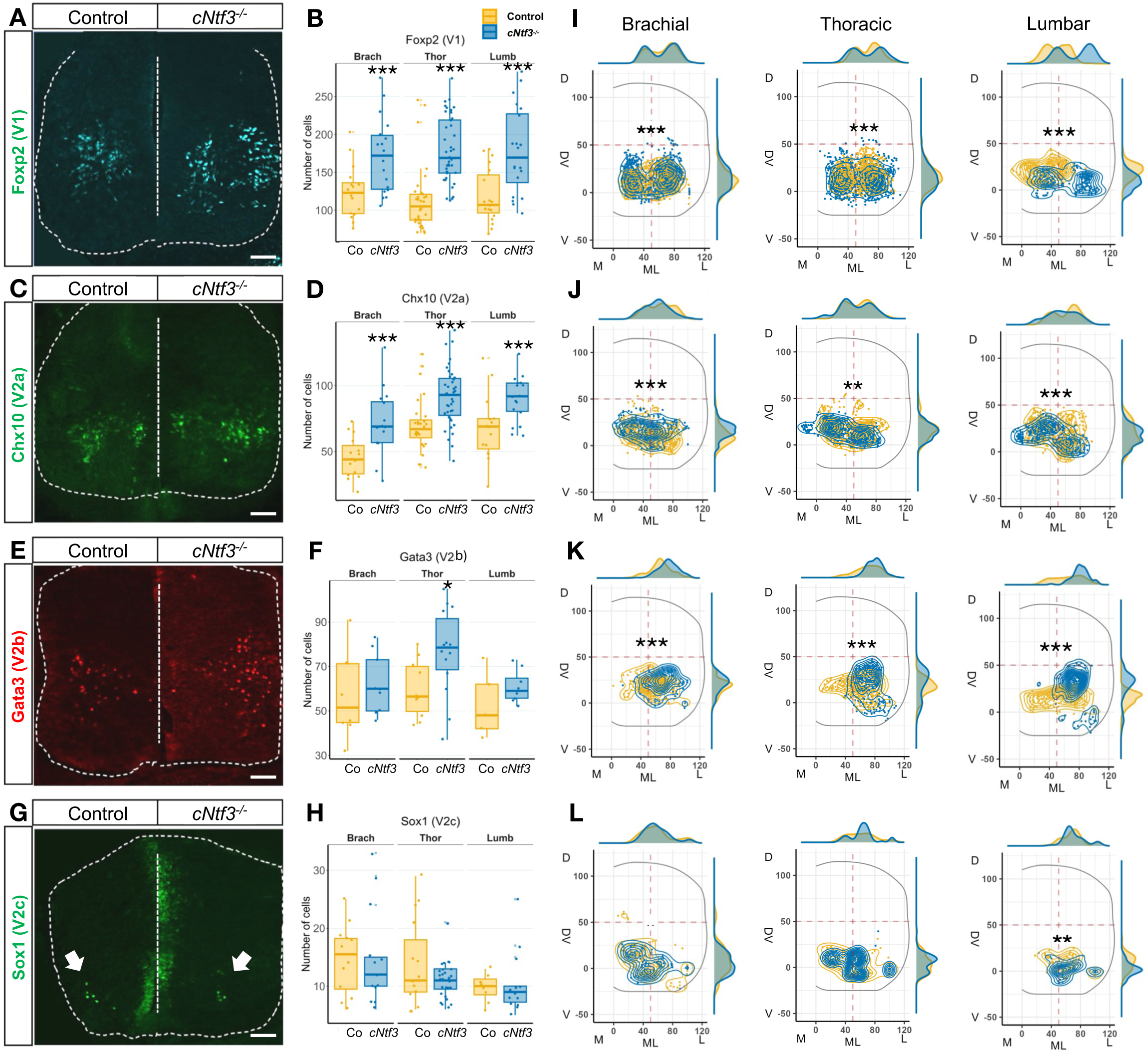
Spinal interneuron number and positioning is altered when Ntf3 is inactivated in the spinal motor neurons. **(A-L)** In MN conditional *Ntf3* mutant embryos (*cNtf3*) at e12.5, the number of V1 (Foxp2) and of V2a (Chx10) INs was significantly increased at all levels of the spinal cord, while V2b (Gata3) only expanded at thoracic level, and V2c (Sox1, white arrows) were not altered. Furthermore, the positioning of all these IN populations on the transversal plane of the spinal cord was altered at all levels of the spinal cord. Labelings are shown at thoracic levels of the spinal cord. Scale bars=100μm. ML: medio-lateral axis; DV: dorso-ventral axis. n=3; *=adjusted p value < 0.05; **=adjusted p value < 0.01; ***=adjusted p value < 0.001

However, IN distribution was additionally affected in the MN-specific OC conditional mutants (Figure 2). Therefore, we also assessed their localization at e12.5 in the absence of Ntf3. At e12.5, V1 INs, which were more numerous (Figure 4A-B), located more laterally and more ventrally than in control embryos, particularly at thoracic and lumbar levels (Figure 4I). This abnormal lateral positioning was maintained at e14.5 (Supplemental Figure S6E). V2a INs remained in a more medio-dorsal position at brachial levels but clustered in the center of the V2a distribution domain at thoracic and lumbar levels at e12.5 (Figure 4J). At e14.5 at brachial levels, V2a did not migrate dorsally as observed in control embryos and located more laterally at brachial levels (Supplemental Figure S6F). In contrast, V2b INs clustered in the lateral half of the medio-lateral axis and located more dorsal than in control embryos at all levels of the spinal cord at e12.5, with some cells aberrantly migrating in a peripheral ventro-lateral position (Figure 4K). This abnormal positioning was maintained at brachial levels at e14.5 (Supplemental Figure S6G). Finally, V2c INs gathered in small, separated groups of cells in the absence of Ntf3, both at e12.5 (Figure 4L) and e14.5 (Supplemental Figure S6H). Thus, in addition to their differentiation, Ntf3 produced by the MNs control the distribution of spinal pre-motor INs in the developing spinal cord, suggesting that Ntf3 coordinates the development of neuronal populations involved in spinal motor circuit formation. To approach this question, we conducted rotarod and balance beam tests to assess coordination in the locomotor circuits built in the absence of Ntf3 from MNs. Both female and male mice deficient for Ntf3 were almost unable to cope with rotarod acceleration, and motor learning was almost absent in these mice, as compared to control littermates (Figure 5A,D). In the balance beam test, females lacking Ntf3 in spinal MNs showed an increased time to cross the beam and increased number of footslips (Figure 5B-C), which was not the case in males (Figure 5E-F). However, learning with time normalized both parameters in females after 3 weeks, as compared to controls (Figure 5B-C). These observations are consistent with the hypothesis that Ntf3 produced by MNs is required to generate locomotor circuits with properly coordinated activity.

**Figure 5.**
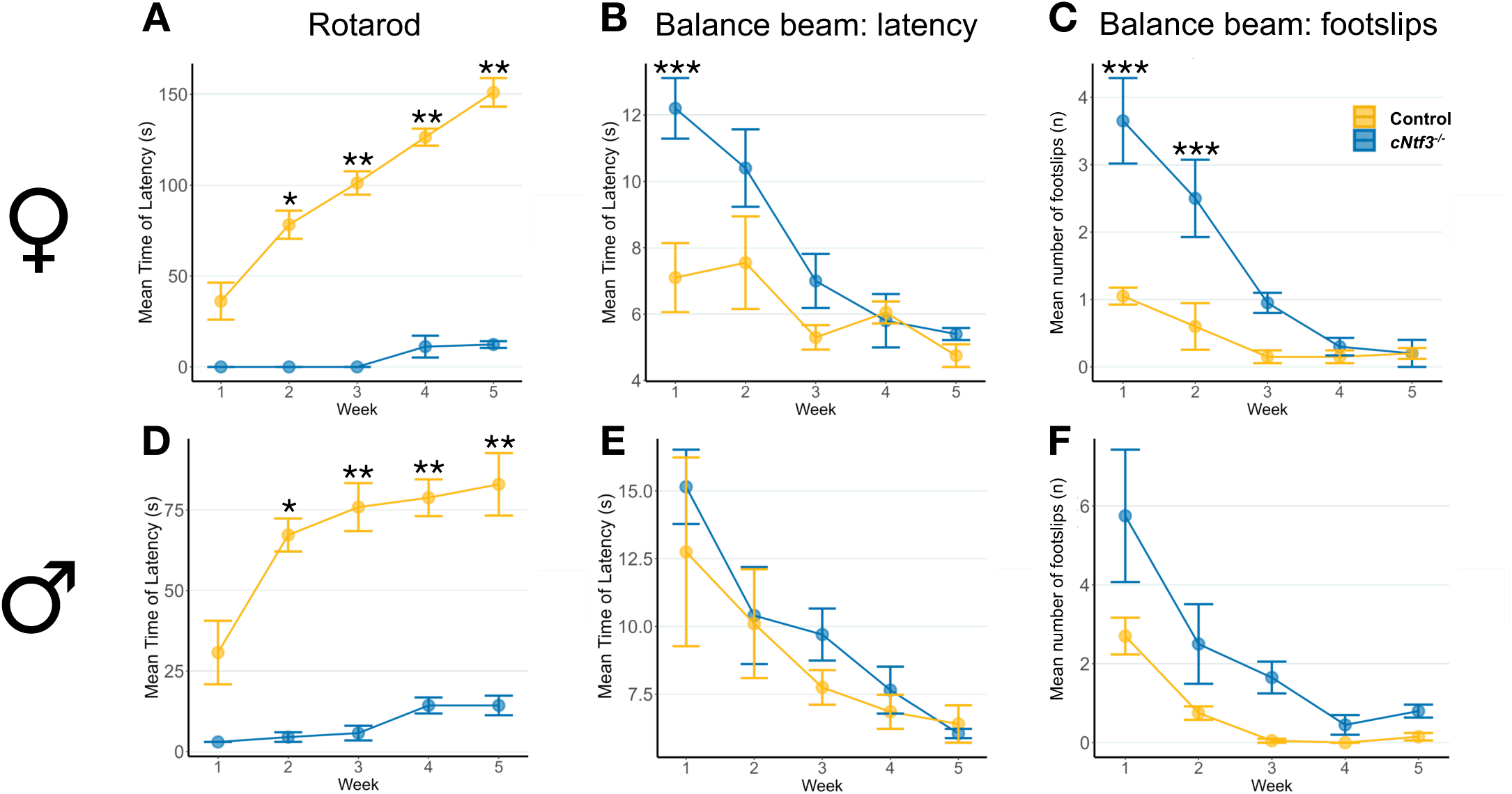
Locomotor coordination is altered when Ntf3 is inactivated in the spinal motor neurons. **(A-C)** In female mice, while control animals demonstrated proper motor coordination and efficient motor learning evidenced by the increase in rotarod latency with time, animals carrying a conditional *Ntf3* mutation in MNs (*cNtf3*) were almost unable to cope with rotarod acceleration and initially showed increased balance beam latency and footslips, which however normalized with time. **(D-F)** In male mice, conditional *Ntf3* mutants showed motor coordination and learning defects on the rotarod but didn’t demonstrate any alteration in the balance beam test. n=8 in each group; *= p value < 0.05; **= p value < 0.01; ***= p value < 0.001

## DISCUSSION

Neural circuits are composed of different types of neurons that interconnect to produce a properly balanced and tightly regulated activity. Cell autonomous mechanisms that control neuronal diversity and connectivity have been widely studied. In contrast, non-cell autonomous processes that coordinate the development of different neuronal types and their integration into functional networks have been less extensively investigated, especially in the spinal cord. Here, we provide evidence that Ntf3 produced in spinal MNs under the control of OC transcription factors coordinate the development of pre-motor INs and the formation of the spinal locomotor circuits.

Our observations in different experimental models highlight the contribution of MNs to IN differentiation and distribution. However, these different models are difficult to compare. MN ablation in *Olig2|DTA* embryos is incomplete, suggesting that residual Ntf3 remains secreted in the ventral spinal cord. *OC* inactivation increases *Ntf3* expression, but to what extent and where downstream signaling is increased remains experimentally out of reach. Furthermore, we obtained evidence that OC factors can travel between cells in the developing spinal cord (data not shown), as previously described for other homeoproteins (reviewed in (Di Nardo et al., 2020)), suggesting that multiple non-cell autonomous processes are at play. Yet, our observations show that disturbing Ntf3 production in spinal MNs impact pre-motor IN development.

During embryonic development, OC transcription factors are present in multiple spinal neuronal populations, including MNs (Francius and Clotman, 2010; Roy et al., 2012), and ventral (Francius et al., 2013; Harris et al., 2019; Stam et al., 2012) or dorsal (Kabayiza et al., 2017) INs. In this tissue, they regulate neuronal differentiation (Harris et al., 2019; Kabayiza et al., 2017; Roy et al., 2012), maintenance (Stam et al., 2012) and IN distribution (Harris et al., 2019; Kabayiza et al., 2017). Although some of these functions are very likely, at least partly, cell autonomous, involving the regulation of *Isl1* in MNs (Rhee et al., 2016; Roy et al., 2012; Toch and Clotman, 2019; Velasco et al., 2017) and of *Pou2f2* or *Nkx6.2* in INs and MNs (Harris et al., 2019; Masgutova et al., 2019; Toch et al., 2020), observations in the constitutive OC mutant embryos suggested that these factors may additionally regulate spinal IN development in a non-cell autonomous manner. Non-cell autonomous regulation of neuronal development in the embryonic spinal cord has already been reported. Murine MNs exert negative feedback on the entry of their own progenitors into differentiation (Park et al., 2013; Sabharwal et al., 2011) and seem to modulate the development of V1 INs (Pfaff et al., 1996), whereas zebrafish primary MNs determine the GABAergic phenotype of Kolmer–Agduhr INs (Seredick et al., 2014). Given that the onset of spinal MN development precedes that of INs, that MNs exert non cell-autonomous regulation of the differentiation of surrounding cells (Park et al., 2013; Pfaff et al., 1996; Sabharwal et al., 2011), that IN alterations have been reported in embryos wherein MN differentiation is strongly affected by the absence of Isl1 (Pfaff et al., 1996), and that OC proteins are critical for proper differentiation of the spinal MNs and for the expression of *Isl1* in these cells (Roy et al., 2012), we hypothesized that OC factors may regulate the observed non-cell autonomous mechanism by acting in spinal MNs. Our observations suggest that MNs control the differentiation and the positioning of the spinal pre-motor INs.

Severe immunotoxic ablation of MNs in an organotypic culture model of embryonic rat spinal cord resulted in a decreased number of inhibitory INs, which was rescued by Ntf3. In the same model, anti-serum blockade of Ntf3 activity was sufficient to more severely deplete inhibitory INs, partly through apoptosis (Bechade et al., 2002). In contrast, conditional *Ntf3* deletion in the mouse rather increased the number of V1 or V2b inhibitory INs, and of V2a excitatory neurons. Critical differences between our *in vivo* model and the highly apoptotic-prone organotypic culture model likely contribute to these discrepancies. Nevertheless, these observations support the contribution of spinal MNs to the control of IN development and the implication of Ntf3 in this process. Opposite results were obtained by other researchers using mouse models lacking almost all MNs due to the elimination of skeletal muscles at late stages of embryonic development (E18.5), where the number of spinal INs remained unaffected (Grieshammer et al., 1998; Kablar and Rudnicki, 1999). This suggests that a critical window of *Ntf3* expression at early stages of spinal development could be critical for proper production of spinal INs. Increased production of specific IN populations in the absence of Ntf3 from MNs, combined with the decrease in V2c observed in the OC mutant embryos wherein *Ntf3* expression is increased, suggest that Ntf3 does not regulate the survival of spinal pre-motor INs, in contrast to sensory neurons (ElShamy and Ernfors, 1996a, b; Liebl et al., 1997), but could rather directly contribute to their differentiation. Consistently, Ntf3 has been shown to cooperate with the morphogen Shh to promote neuronal differentiation (Dutton et al., 1999). In addition, our observations suggest that Ntf3 produced by spinal MNs regulate the migration of pre-motor INs, as the distribution of these cells was affected in embryos lacking Ntf3 from MNs. The characterization of functionally distinct IN subpopulations unveiled a strong correlation between the distribution of each IN subset and their contribution to distinct microcircuit modules (Bikoff et al., 2016; Borowska et al., 2013; Goetz et al., 2015; Hayashi et al., 2018; Hilde et al., 2016; Tripodi et al., 2011). These data support a model wherein correct localization of spinal IN subsets is critical for proper formation of sensory-motor circuits, highlighting the importance of a strict regulation of short-distance neuronal migration in the developing spinal cord. However, genetic determinants that control spinal IN migration have only been sparsely identified (Blacklaws et al., 2015; Harris et al., 2019; Hilde et al., 2016; Kabayiza et al., 2017; Masgutova et al., 2019). Furthermore, non-cell autonomous mechanisms contribute to this control. Indeed, commissural axons have been shown to preconfigure ventral IN cell body position in the contra-lateral side of the spinal cord, acting either as border landmarks or as cellular guides for migration (Laumonnerie et al., 2015). This highlights a complex interplay between neuronal identity and neuronal positioning, jointly controlled by cell autonomous and non-cell autonomous mechanisms. Given that IN differentiation and IN distribution are affected upon *Ntf3* deletion from MNs, separate contribution to one or the other process remains to be investigated. Transcriptomic characterization of alterations in pre-motor INs upon changes in Ntf3 production should address this question.

Ntf3 has been implicated in non-cell autonomous processes in other parts of the central nervous system. For example, in the developing cortex, Ntf3 produced by postmitotic neurons modulates the fate of progenitor cells during radial expansion (Parthasarathy et al., 2014). In the avian inner ear, it is required for proper development of the intrinsic properties of low-frequency neurons of the tonotopic axis in the cochlea (Takahashi and Sanchez, 2020), and could have a role in the postnatal inner ear activity, where it is transported by cochlear axons into the ventral cochlear nucleus (Feng et al., 2010). Furthermore, work in the chicken retina demonstrated that retinal ganglion cells can produce Ntf3 and transport it towards the optic tectum to control the survival and connectivity of superficial tectum cells (von Bartheld and Butowt, 2000). Importantly, spinal MNs have been shown to be necessary for survival and for peripheral or central projections of sensory neurons, and Ntf3 produced by the MNs partly contributes to survival and central projections of proprioceptive neurons, likely through transaxonal signaling (Usui et al., 2012). Therefore, the motor defects we observed in the MN-specific *Ntf3* mutants likely result from combined alterations of proprioceptive inputs and of the intrinsic organization and activity of spinal motor circuits. Nevertheless, our differentiation and distribution studies demonstrate a direct impact of Ntf3 produced in MNs on the development of spinal INs. Thus, Ntf3 contributes to regulate neuronal differentiation, survival, connectivity and activity in multiple neural structures.

At early stages of spinal cord development, *Ntf3* expression was restricted to MNs and its cognate receptor TrkC was detected in surrounding IN populations, consistent with previous reports (Buck et al., 2000; Henderson et al., 1993; Usui et al., 2012). OC factors moderate *Ntf3* expression levels in MNs, as previously observed for *Nkx6.2* (Toch et al., 2020) or for *Pou2f2* in ventral and in dorsal IN populations (Harris et al., 2019; Masgutova et al., 2019), suggesting that Ntf3 production in MNs must remain moderate to ensure proper development of pre-motor INs. Increased Ntf3 production has previously been shown to disrupt nervous system development or activity. *Ntf3* overexpression alters neuronal differentiation in the neocortex (Parthasarathy et al., 2014). In inner ear supporting cells or hair cells, it increased ribbon synapse density in postnatal cochlea and reduced auditory brainstem response thresholds at high frequencies (Wan et al., 2014). Increased production in the dorsal amygdala resulted in reduced anxious temperament and altered function of corresponding neural circuit (Fox et al., 2019). Thus, controlled levels of Ntf3 are critical for proper development and homeostasis of the nervous system.

Beside passive diffusion of signaling molecules in the extracellular space or transaxonal signaling as observed for Ntf3 (Usui et al., 2012), alternative mechanisms could additionally contribute to non-cell autonomous regulation of CNS development. Extracellular vesicles, which are also produced in the developing human spinal cord (Cau et al., 2022), have recently been shown to regulate the dorso-ventral differentiation and distribution of cortical INs. Extracellular vesicles produced in the ventral regions of the developing cortex contain secreted molecules, membrane proteins and ligand/receptor partners, and modulate fate and migration of ventrally-produced cortical INs (Pipicelli et al., 2023). Even more surprisingly, tunnelling nanotubes able to transfer molecules, endosomes and even mitochondria have recently been observed in living zebrafish embryos (Korenkova et al., 2025). Whether these transfer modalities are also in action in the mouse developing spinal cord remains to be assessed.

## MATERIALS AND METHODS

### Ethic statement and mouse lines

All experiments were performed strictly in accordance with the European directive 2010/63/UE. Mice were maintained in a conventional facility and fed in standard conditions (mice maintenance and mice breeding diets, Carfil Quality, Belgium), on a 14 h light/10 h dark cycle. Food and water were available *ad libitum.* Experimental procedures on animals were approved by the animal ethics committees of the Université catholique de Louvain (Permit Numbers: 2013/UCL/MD/11; 2017/UCL/MD/008; 222801). The mutant strain mice were crossed and the morning the vaginal plug was detected was defined as embryonic day (e) 0.5. A minimum of three embryos (n≥3) of the same genotype was analyzed in each experiment. The embryos were harvested at embryonic days (e)10.5, e11.5, e12.5 or e14.5 depending on the mouse line. Constitutive Oc1^+/-^;Oc2^+/-^ mutants were crossed to obtain Oc1^-/-^;Oc2^-/-^ double-knockout embryos (Clotman et al., 2005). *Olig2|DTA* embryos, wherein the *Olig2* promoter drives expression of the diphteria toxin subunit A (DTA) in MN progenitors, have been obtained by crossing *Olig2-Cre* mice (Dessaud et al., 2007) with *Rosa26-flSTOP-DTA* mice (Ohnmacht et al., 2009). The Oc1^flox/flox^; Oc2^flox/flox^ mutant mice were crossed with Rosa26R- YFP/Olig2-Cre transgenic mice bearing heterozygous-null mutations for Oc1 and Oc2 genes (Rosa26-YFP;Olig2Cre;Oc1^+/−^;Oc2^+/−^) to obtain conditional double inactivation of Oc1 and Oc2 in MNs (Toch et al., 2020). The combined inactivation of Oc1 and Oc2 in MNs completely abolishes the expression of *Oc3* in the developing spinal cord (Kabayiza et al., 2017; Roy et al., 2012). Ntf3^+/flox^ mice (B6.129 S4-Ntf3tm2Jae/J, The Jackson Laboratory) were crossed first with PGK-Cre mice to obtain a transgenic line with a constitutive heterozygous inactivation of *Ntf3* (Ntf3^+/-^), then with the Olig2-Cre mice (Olig2Cre;Ntf3^+/-^). The Ntf3^flox/flox^ mutant mice were crossed with the Olig2Cre;Ntf3^+/-^ mice to obtain a conditional inactivation of *Ntf3* in MNs. PCR genotyping protocols and primers are available upon request.

### *In situ* hybridization and immunofluorescence

For *in situ* hybridization (ISH), collected embryos were fixed in ice-cold 4% paraformaldehyde (PFA) in phosphate buffered saline (PBS) overnight at 4 °C, washed thrice in PBS for 10 minutes, incubated in PBS/30% sucrose solution overnight at 4 °C, and embedded and frozen in PBS/15% sucrose/7.5% gelatin. Sixteen micrometer sections were prepared, and ISH was performed as previously described (Beguin et al., 2013; Francius et al., 2016; Pelosi et al., 2014) with DIG-conjugated Ntf3 antisense RNA probes (primer pair: 5ʹ-GATCCAGGCGGATATCTTGA-3ʹ and 5ʹ-CGGACATAGGTTTGCGAAGT-3ʹ). Control or Ntf3 conditional mutant sections were placed adjacent on the same histology slides to minimize inter-slide variations of ISH signals.

For immunofluorescence, collected embryos were immersion-fixed in 4% PFA/PBS for 15, 25 or 35 minutes at 4°C according to their embryonic stage, and processed as for ISH. Immunolabeling was performed on 14 µm serial cryosections as previously described (Debrulle et al., 2020; Francius et al., 2013). Primary antibodies against the following proteins were used: sheep anti-Chx10 (Exalpha Biologicals #X1179P) at 1:200 or guinea pig anti-Chx10 (Peng et al., 2007) at 1:3000, rabbit anti-cleaved Caspase-3 (Asp175) antibody (Cell Signaling #9664) at 1:1000, mouse anti-Evx1 (DSHB #99.1-3A2) at 1:2000 or rabbit anti-Evx1 (Moran-Rivard et al., 2001) at 1:300, rabbit anti-Foxd3 at 1:5000 or guinea pig anti-Foxd3 (Muller et al., 2005) at 1:5000, goat anti-Foxp2 (Abcam #ab1307) at 1:2000, guinea pig anti-Gata2 (Peng et al., 2007) at 1:3000, rat anti-Gata3 (Absea Biotechnology #111214D02) at 1:15 or rabbit anti-Gata3 (Cell signaling #5852) at 1:200, guinea pig anti-MafA (Gierl et al., 2006) at 1:500, goat anti-Isl1 (R&D #AF1837) at 1:1000, rabbit anti-MafA (Novus Biological #NB400-137) at 1:500, mouse anti-MNR2 (DSHB #81.5C10) at 1:1000, Mouse anti-Nkx2.2 (DSHB #74.5A5) at 1:20, rabbit anti-Otp at 1:1600 (ThermoFisher #PA5-89060), rabbit anti-Prdm8 at 1 :200 or mouse anti-Prdm8 at 1:200(Komai et al., 2009), goat anti-Sox1 (Santa Cruz #sc-17318) at 1:500 or rabbit anti-Sox1 (Panayi et al., 2010) at 1:400 and goat anti-TrkC (R&D Systems #AF1404) at 1:500. The secondary antibodies used are donkey anti-chicken/Alexafluor 488, donkey anti-goat/AlexaFluor 594, donkey anti-goat/AlexaFluor 488, donkey anti-goat/AlexaFluor 647, donkey anti-mouse/AlexaFluor 594, donkey anti-mouse/AlexaFluor 488, donkey anti-rabbit/AlexaFluor 594, donkey anti-rabbit/AlexaFluor 488, donkey anti-rabbit/AlexaFluor 647, donkey anti-rat/AlexaFluor 594, donkey anti-rat/AlexaFluor 488, donkey anti-guinea pig/Alexa fluor 488 purchased from ThermoFisher Scientific or Jackson Laboratories and used at 1:500 to 1:2000 dilutions.

Immunofluorescence and ISH images from cryosections were acquired on an EVOS FL Auto Imaging System (ThermoFisher Scientific), a Zeiss AxioSkop2 or Zeiss AXIO Observer Z1 epifluorescence microscopes. The images were processed using Adobe Photoshop CS5 or ImageJ (Fiji) softwares to match brightness and contrast. The rostrocaudal levels of the spinal cord were precisely aligned using the distribution of Foxp1 in the Lateral Motor Columns (LMCs) at brachial or lumbar levels of the spinal cord (Rousso et al., 2008; Roy et al., 2012).

### *In ovo* electroporation

The *in ovo* electroporations of chicken embryonic spinal cord were performed at stage HH12 (around forty-eight hours of development), and embryos were collected 24, 48 or 72 h after electroporation. The coding sequence of *Ntf3* (primer pair: 5’-CGGATGCCATGGTTACTTCT-3’ and 5’-TGCCAATTCATGTTCTTCCA-3’) was cloned into the vector pHb9-MCS-IRES-EGFP to increase the expression of the gene in the MNs. pHb9-Ntf3-IRES-GFP vector (2 µg/µl) was electroporated using the EGFP signal to visualize cells expressing *Ntf3*. However, given the transient expression of such constructs after electroporation, maximal GFP production could only be visualized 24h after electroporation (data not shown). The collected embryos were fixed in ice-cold PBS/4% PFA for 15, 20 or 25 minutes, depending on their stage, and processed as described above for ISH or immunofluorescence. The MNs and ventral IN populations were analyzed using 5-10 sections per embryo, the contralateral side of the spinal cord being considered as a perfect stage- and experimental condition-matching control. The labelled cells were counted on both sides of the spinal cord, with the counting tool of Image J (Fiji).

### FACS, RNA purification and RNA-sequencing

Spinal cords from e10.5 control or Rosa26-YFP;Olig2Cre;Oc1^Δ/−^;Oc2 ^Δ /−^ embryos were harvested and dissociated using a neural tissue dissociation kit (MACS; Miltenyi Biotec #130-092-628), dissociated cells were sorted by FACS (BD FACSAria III) to collect YFP-positive cells. The sorted cells were collected in TRIzol reagent, and RNA was purifed with the Rneasy micro kit (QIAGEN #74004). RNA concentration and quality were assessed using a Bioanalyzer (Agilent) and submitted to Genewiz to prepare an ultra-low input RNA-seq library before sequencing with an Illumina HiSeq. Preliminary data were analyzed by Genewiz using the standard RNA-seq data analysis package. RNAseq data have been deposited in the GEO repository (accession number: GSE141949)(Toch et al., 2020).

### Motor behavior tests in adult mice

Behavior tests were performed on male (n=8) or female (n=8) mice, comparing control Olig2Cre;Ntf3^+/-^ mice with Olig2Cre;Ntf3 ^Δ/-^ conditional mutant mice which lack Ntf3 in MNs, starting at two months of age. The rotarod and the balance beam tests were performed as previously described (Audouard et al., 2012; Renaux et al., 2024).

### Statistical analyses

For each of the embryos analyzed (n≥3), neuron quantifications were performed using the count analysis tool of ImageJ on sections of the spinal cord at brachial, thoracic or lumbar levels, except for chicken embryos that are too early to delineate different levels in the spinal cord. Graphs were generated using R. The statistical tests applied to compare the cell count comparing two groups (controls versus constitutive or conditional mutants) was performed in R [R Core Team 2024] (version 4.4.2) either with a mixed model (using the R packages lme4 (Bates et al., 2015) version 1.1-37 and lmerTest (Kuznetsova et al., 2017) version 3.1- 3) involving a random embryo effect if the random effect was significant or a linear regression if it was deemed non-significant. A correction for multiple testing with the False Discovery Rate procedure was applied to the obtained p-values.

The spatial distribution of the motor neurons and ventral IN populations was analyzed using a gradient map. In a transverse section of the spinal cord, the height (H) was defined as the distance from the ventral limit of the central canal to the most dorsal edge of the spinal cord, and the width (W) was the distance from the central canal to the lateral edge. For each population, several sections were analyzed at brachial, thoracic and lumbar levels. The distance (dIN) and angle (αIN) were measured from the ventral limit of the central canal to the cell soma using the ruler analysis tool in Image J. The dorso-ventral (DV) and medio-lateral (ML) positions of the cells were expressed as percentages of spinal cord height and hemicord width respectively: DV position and ML position were defined as (dIN ∗ sin αIN)/H and (dIN ∗ cos αIN)/W, respectively, and ML versus DV values were plotted using R with a 2D kernel density (Harris et al., 2019; Kabayiza et al., 2017). For each section and level, a non parametric multivariate ANOVA-type statistic was performed (R package npmv, version 2.4 (Burchett et al., 2017)) to assess if the locations of DV and ML in the two conditions are different.

For the balance beam and rotarod tests, differences between the two experimental groups were evaluated with the JMP software using either a Wilcoxon-Mann-Whitney’s non parametrical test, a Welch’s *t*-test or a Student’s *t*-test, depending on the normality and the homoscedasticity of the data. In all statistical analyses, a value of p < 0.05 was defined as significant.

The complete statistical analysis is available here: https://github.com/ManonMartin/Non-cell_autonomous_regulation_of_spinal_motor_circuit_development_Angla-Navarro-Dominguez-Bajo

## ACKNOWLEDGMENTS

We thank previous members of the NEDI lab and members of the AMCB group for material, technical support and discussions, Coralie Piget at the “Animalerie Carnoy” (ANCA) facility for expert assistance in mouse colony care and management, and Catherine Rasse for expert input on quantifications and statistical analyses. We are indebted to Pr. Yoichi Shinkai, Dr. Stavros Malas, Dr. Martyn Goulding, Dr. Kamal Sharma, Dr. Sam Pfaff, Dr. Thomas Müller, Dr. Frank Grosveld, Dr. Yoichi Shinkai, Dr. Sarah E. Ross and Dr. Stavros Malas for antibodies and reagents, and to Dr. Andrea B. Huber for the generous gift of *Olig2|DTA* embryo sections. The anti-Evx1, -MNR2 and -Nkx2.2 antibodies were obtained from the Developmental Studies Hybridoma Bank, created by the NICHD of the NIH and maintained at The University of Iowa, Department of Biology, Iowa City, IA 52242. Research in the FC group is supported by grants from the « Fonds spéciaux de recherche » (FSR) of the Université catholique de Louvain, by the “Actions de Recherche Concertées (ARC)” #17/22-079 of the “Direction générale de l’Enseignement non obligatoire et de la Recherche scientifique – Direction de la Recherche scientifique – Communauté française de Belgique” and granted by the “Académie universitaire ‘Louvain’”, by a “Projet de recherche (PDR)” #T.0117.13 and an “Equipement (EQP)” funding #U.N027.14 of the Fonds de la Recherche Scientifique (F.R.S.-FNRS) and by the Association Belge contre les Maladies neuro-Musculaires (ABMM). Work in XM lab is supported by grants from the National Eye Institute of the National Institutes of Health (R01EY020545 and R01EY029705). AAN held a PhD grant from the F.R.S.-FNRS, Belgium; MT held a specialization grant from the FRIA (F.R.S.-FNRS); JZ holds a PhD grant from the China Scholarship Council (CSC, No. 202208440068) and a UCLouvain-CSC Ph.D. co-funding fellowship from UCLouvain; MF holds a PhD grant from the Fonds Spéciaux de Recherche (FSR) from UClouvain; ADB and MHF held a postdoctoral researcher grant from the F.R.S.-FNRS. FC is a Research Director of the F.R.S.-FNRS.

## AUTHOR CONTRIBUTION

AAN: Conceptualization, Investigation, Data analysis, Data interpretation, Writing - original draft

ADB: Conceptualization, Investigation, Data analysis, Data interpretation, Writing – original draft

MT: Conceptualization, Investigation, Data analysis, Data interpretation

CF: Conceptualization, Investigation, Data analysis, Data interpretation

MHF: Conceptualization, Investigation, Data analysis, Data interpretation

JZ: Investigation, Data analysis

MF: Investigation, Data analysis

OS: Conceptualization, Investigation, Data analysis

MM: Data analysis

XM: Provided experimental model

RR: Data interpretation

FG: Data interpretation

FC: Conceptualization, Supervision, Funding acquisition, Data interpretation, Writing – original draft

No competing interests declared

---

**Figure S1.** The MN progenitor domain is not altered in *Olig2|DTA* embryos. At e10.5, the number of MN progenitors characterized by the presence of Olig2 (green) in the absence of Ascl1 (V2 progenitors, red) or Nkx2.2 (V3 progenitors, blue) is similar to control embryos. Scale bar=100μm.

**Figure S2.** Spinal interneuron number and positioning is altered when Onecut factors are absent from the spinal motor neurons. **(A-H)** In MN conditional *Oc1/Oc2* double-mutant embryos (cDKO) at e14.5, the number of V2c INs identified by the presence of Sox1 outside of the ventricular zone was significantly reduced at all levels of the spinal cord. Furthermore, the positioning of the V1 (Foxp2), V2a (Chx10), V2b (Gata3) and V2c INs on the transversal plane of the spinal cord was altered. ML: medio-lateral axis; DV: dorso-ventral axis. n=3; *=adjusted p value < 0.05; **=adjusted p value < 0.01; ***=adjusted p value < 0.001

**Figure S3.** Pathway analysis of down- or up-regulated genes in spinal MN deficient in Onecut factors. Genes showing altered expression levels are enriched in the “cytokine-cytokine receptor interaction” and in the “Axon guidance” pathways. Down-regulated genes are in blue and up-regulated genes in red (see GEO repository, accession number GSE141949 for more details).

**Figure S4.** Ntf3 overexpression after chicken embryonic spinal cord electroporation mildly impacts spinal interneuron development. **(A)** The pHb9-Ntf3-IRES-EGFP expression vector was electroporated in the chicken embryonic spinal cord. **(B)** Immunofluorescence labelings for Isl1 to identify MNs (given its transient expression, maximal GFP production could only be visualized 24h after electroporation) and Chx10 to visualize V2a INs 48h after electroporation. **(C)** Quantification of IN populations in control (Ctrl, contralateral sides) or electroporated (E) sides of the spinal cord. The number of V2a INs was mildly decreased after *Ntf3* overexpression, while other IN populations were not affected. Scale bars=100μm. n=3; *=adjusted p value < 0.05

**Figure S5.** Ntf3 inactivation does not increase spinal cord cell death. Conditional inactivation of *Ntf3* in MNs does not induce cell death in the spinal cord at e12.5, as evidenced by the absence of cleaved-Caspase3. Cell death of sensory neurons in the dorsal root ganglia (arrowheads) served as a positive labeling control.

**Figure S6.** Spinal interneuron number and positioning is altered when Ntf3 is inactivated in the spinal motor neurons. **(A-H)** In MN conditional *Ntf3* mutant embryos (*cNtf3*) at e14.5, the number of V1 (Foxp2) INs was significantly increased at all levels of the spinal cord, while V2a (Chx10) only expanded at lumbar level and V2b (Gata3) or V2c (Sox1, white arrows) were not altered. Furthermore, the positioning of all these IN populations on the transversal plane of the spinal cord was altered at all levels of the spinal cord. ML: medio-lateral axis; DV: dorso-ventral axis. n=3; *=adjusted p value < 0.05; **=adjusted p value < 0.01; ***=adjusted p value < 0.001

