## Supplemental figures for "Neurotrophin-3 produced by motor neurons non-cell autonomously regulates the development of pre-motor interneurons in the developing spinal cord"

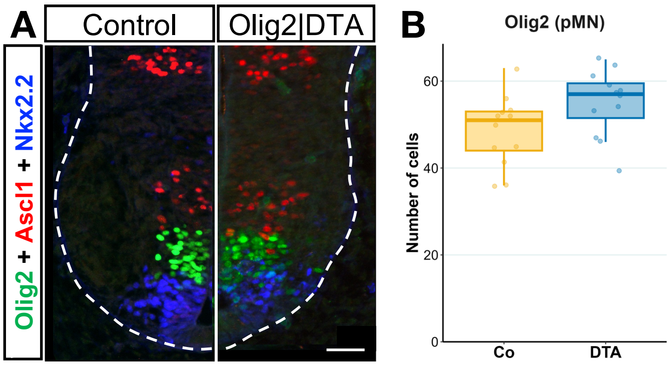


Angla *et al.*, Figure S1

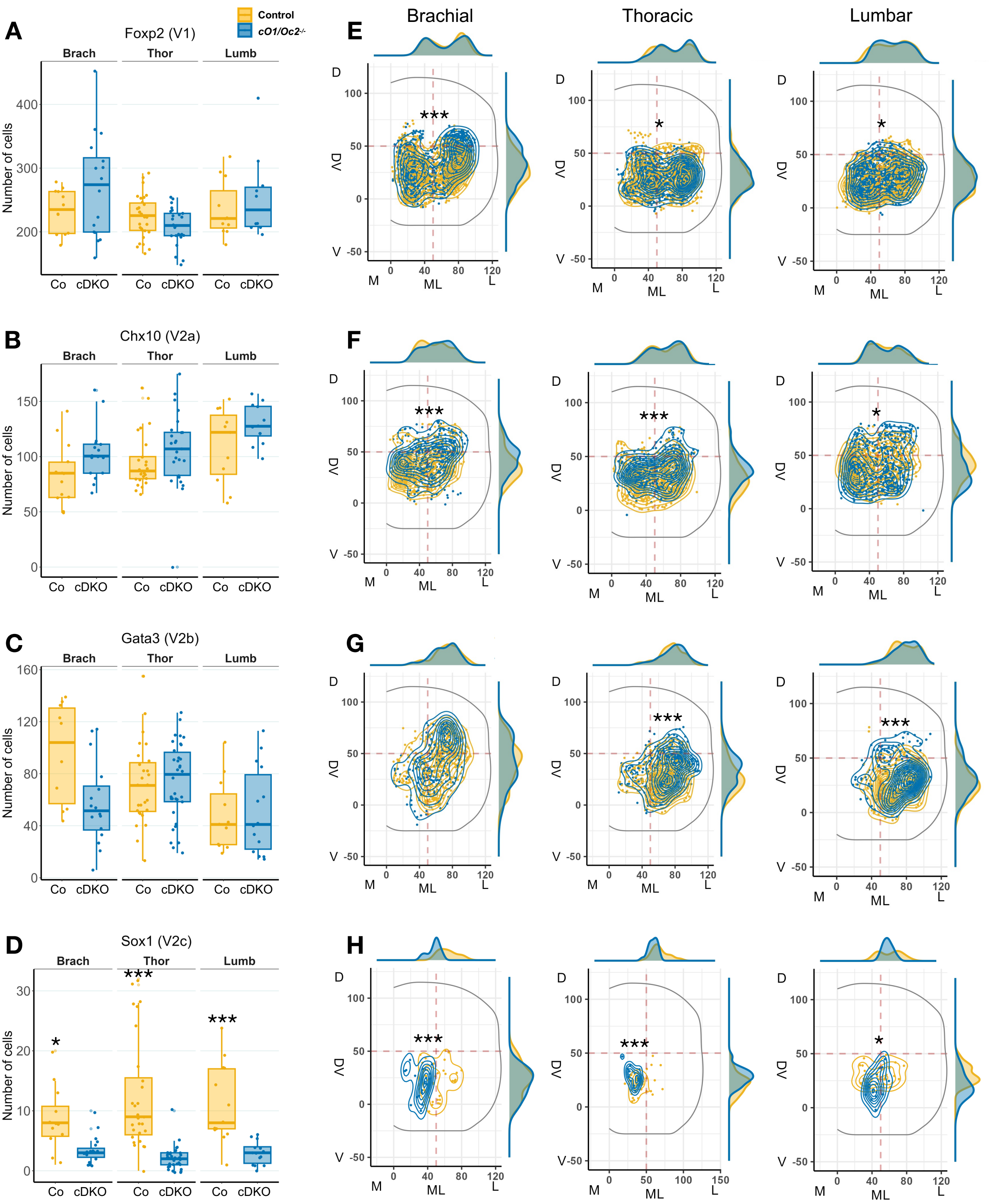


Angla *et al*., Figure S2

### Cytokine-cytokine receptor interaction

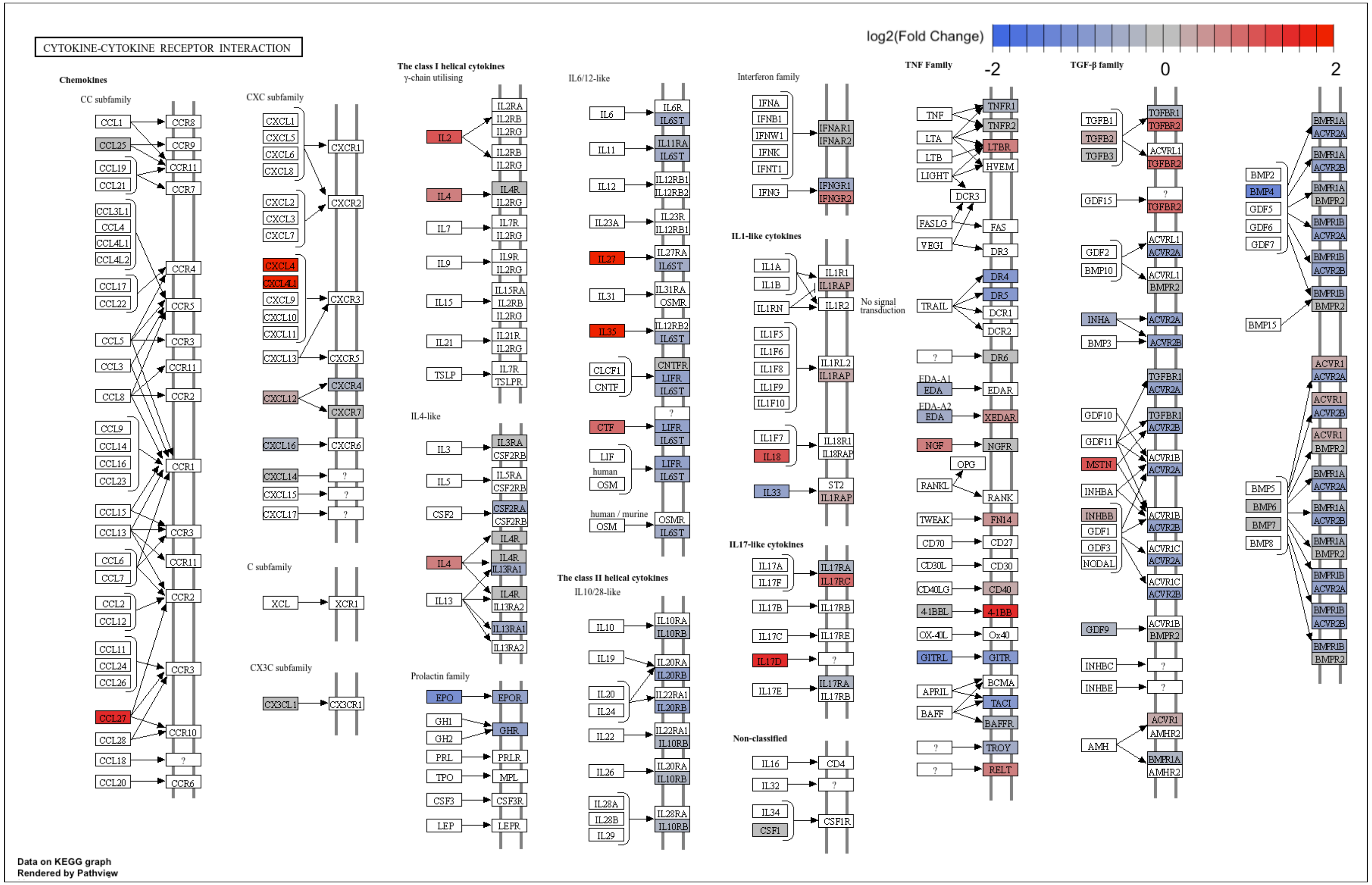


### Axon Guidance

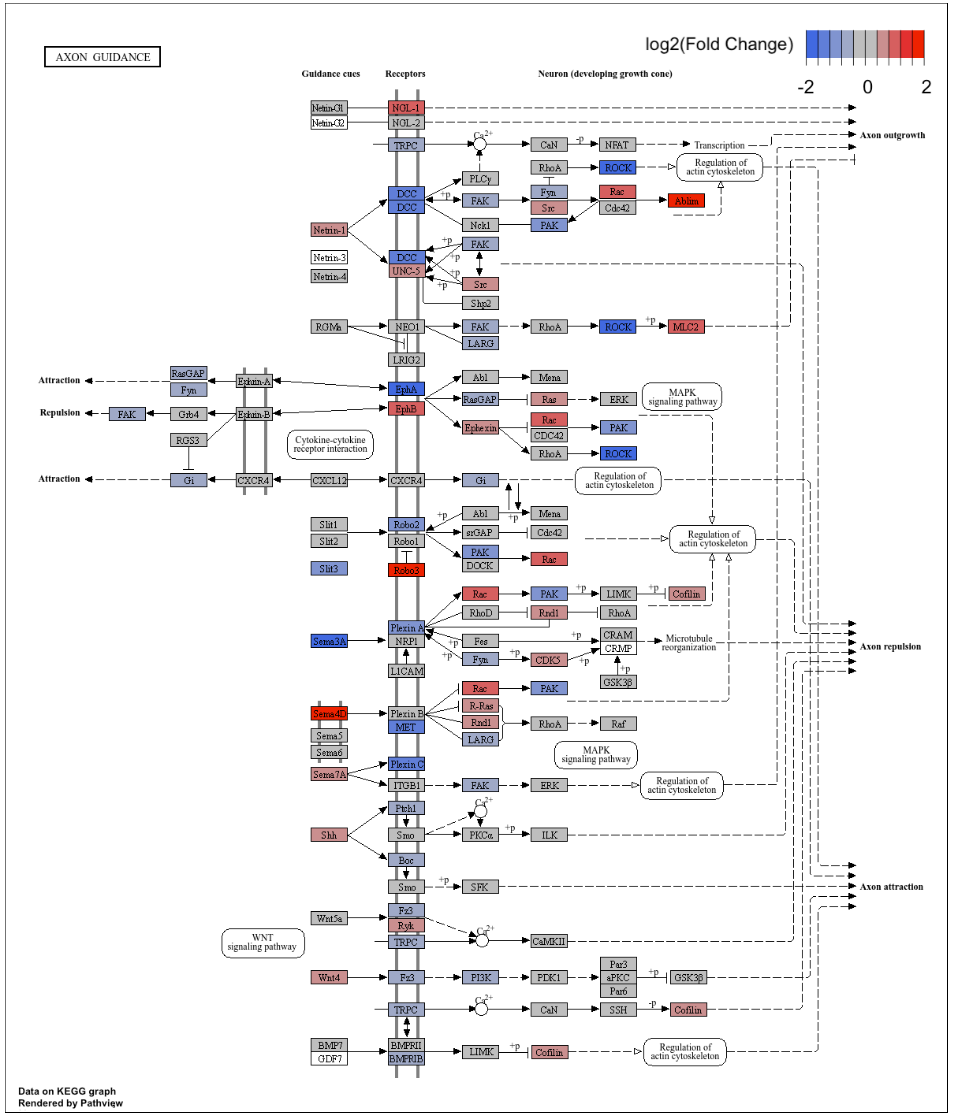


Angla *et al.*, Figure S3

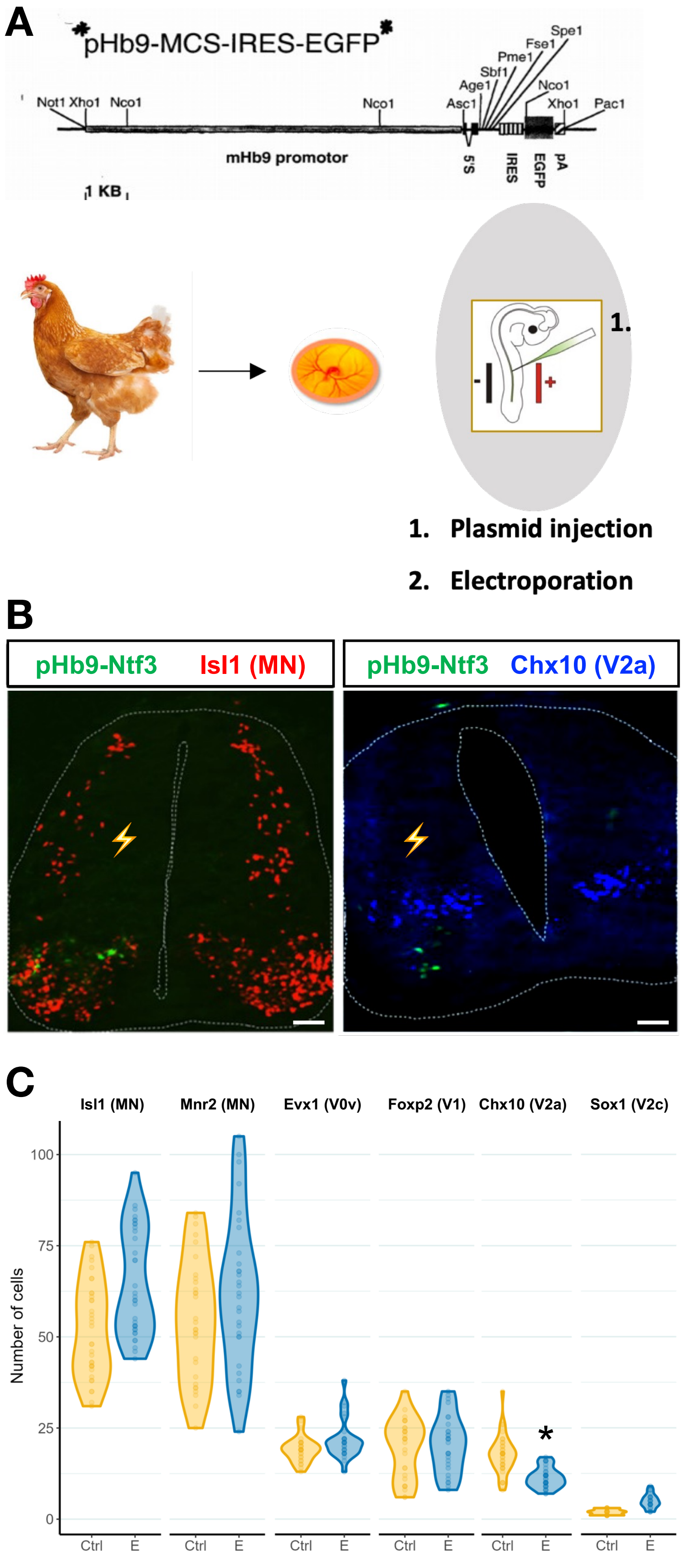


Angla *et al.*, Figure S4

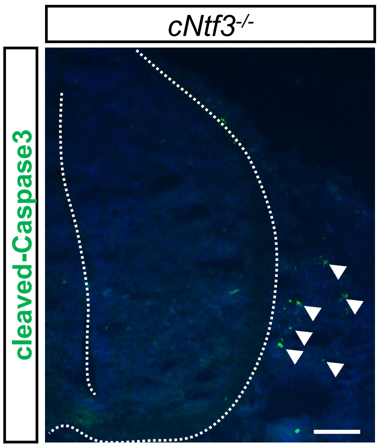


Angla *et al.*, Figure S5

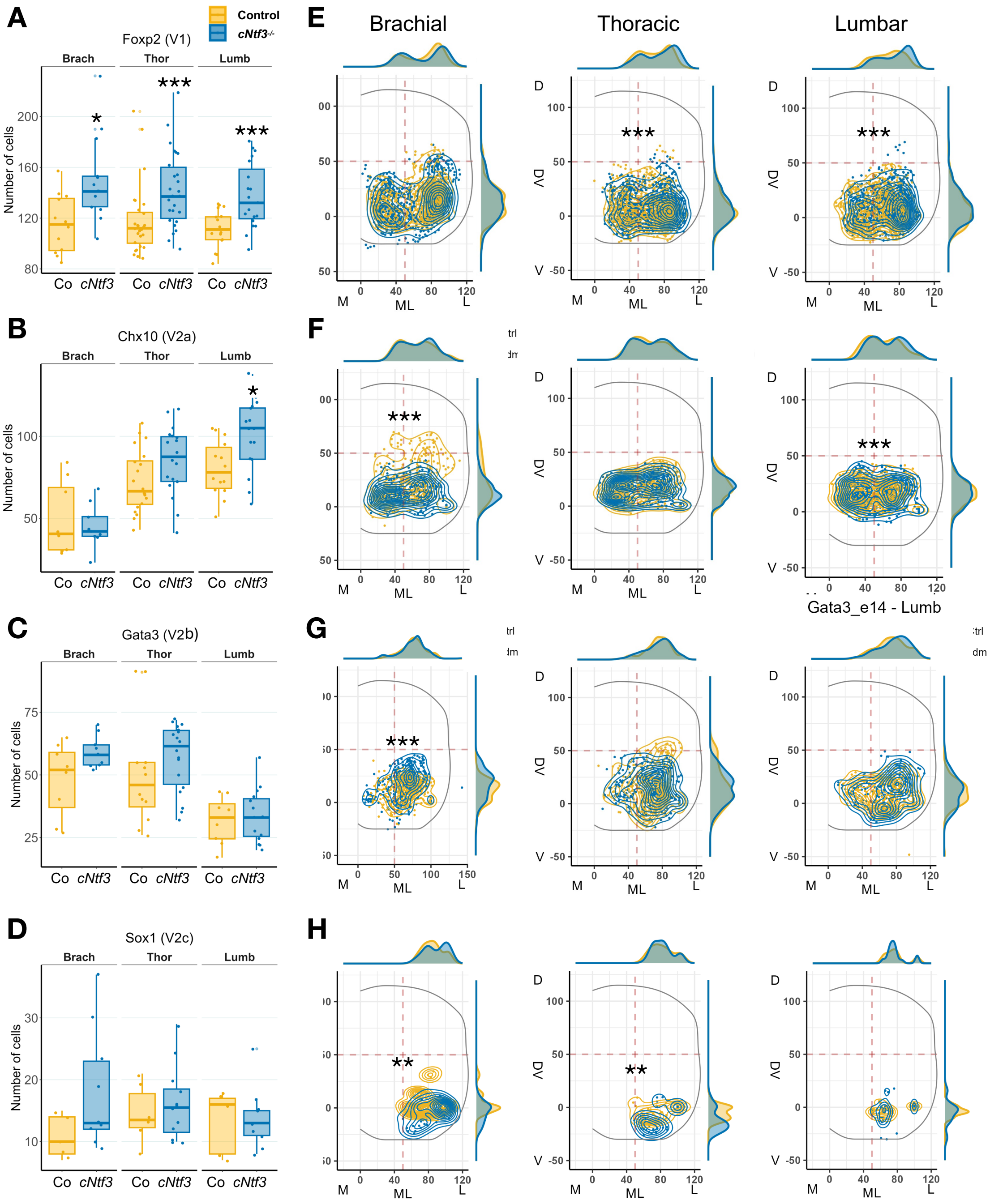


Angla *et al.*, Figure S6
